# Exploring Breast Cancer-Related Biochemical Changes in Circulating Extracellular Vesicles Using Raman Spectroscopy

**DOI:** 10.1101/2024.09.25.614667

**Authors:** Arianna Bonizzi, Lorena Signati, Maria Grimaldi, Marta Truffi, Francesca Piccotti, Stella Gagliardi, Giulia Dotti, Serena Mazzucchelli, Sara Albasini, Roberta Cazzola, Debanjan Bhowmik, Chandrabhas Narayana, Fabio Corsi, Carlo Morasso

**Affiliations:** Department of Biomedical and Clinical Sciences, Università di Milano 20157 Milano, Via Giovanni Battista Grassi, 74, 20157 Milan, Italy; Istituti Clinici Scientifici Maugeri IRCCS, Via Maugeri 4, 27100 Pavia, Italy; Molecular Biology and Transcriptomics Unit, IRCCS Mondino Foundation, Via Mondino 2, 27100 Pavia, Italy; Transdisciplinary Biology Program, Rajiv Gandhi Centre for Biotechnology, Thycaud P.O., Poojappura, Thiruvananthapuram 695014, India; Chemistry and Physics of Materials Unit, Jawaharlal Nehru Centre for Advanced Scientific Research, Jakkur P.O., Bangalore 560064, India

**Keywords:** Breast cancer, Extracellular vesicles, Raman Spectroscopy, Lipoproteins, Biochemical profile

## Abstract

Extracellular vesicles (EVs) are a subgroup of the circulating particles, released by cells in both normal and diseased states, carrying active biomolecules. They have gained significant attention as potential cancer biomarkers, particularly in breast cancer (BC). Previous research showed variations in EVs content and quantity between BC patients and healthy controls (HC). However, studying EVs biochemical profile remains challenging due to their low abundance and complex composition. Additionally, EVs may interact with other plasma components, like lipoproteins (LPs), forming a so called “biomolecular corona” that further complicates their analysis. Here, Raman spectroscopy (RS) is proposed as a fast tool to obtain the biochemical profile of circulating EVs in the context of BC. RS was employed to differentiate various extracellular particles (EPs) in blood, including LPs and EVs. The study also evaluated RS’s capability to quantify major classes of biomolecules and compared these results with those obtained by traditional biochemical assays. Finally, compositional differences in large EVs (lEVs) and small EVs (sEVs) were assessed between 30 HC and 34 BC patients. RS revealed the existence of distinct biochemical signatures associated with BC, highlighting increased levels of nucleic acids and lipids in the BC group.

## 1. Introduction

Breast cancer (BC) is the most common cancer diagnosed in women and and the second leading cause of cancer-related deaths among women worldwide (Menon et al., 2024). The early detection and monitoring are critical factors for the management of BC patients drastically improving their survival rates (Cueva Bañuelos et al., 2020; Zielonke et al., 2020). Finding a method to achieve these objectives without subjecting patients to invasive instrumental investigations, which cause discomfort for them and incur high costs for the healthcare systems, remains complex.

In recent years, liquid biopsy emerged as a non-invasive and promising approach for diagnosing and monitoring BC. Liquid biopsy refers to the detection in blood of characteristic constituents of cancers, such as nucleic acids, including cell-free DNA (cfDNA) or circulating tumor DNA (ctDNA), or circulating tumor cells (CTCs) in a way that allows to detect cancer-related biomolecular features without using invasive tissue biopsies (Boukovala et al., 2024; Wang et al., 2024). Among the various possible components to be analyzed in a liquid biopsy, extracellular vesicles (EVs) have attract considerable attention due to their potential in cancer diagnosis, prognosis, and therapy response monitoring because of their early release from cancer cells (Dorado et al., 2024; Li et al., 2021).

EVs act as messengers in intercellular communication, transporting proteins, lipids, and nucleic acids that can influence various biological processes (Welsh et al., 2024). Several studies have demonstrated that EVs are extensively involved in the major pathways implicated in BC development, proliferation, migration, tumor microenvironment remodeling, immunosuppression and drug resistance (Berumen Sánchez et al., 2021; Qian et al., 2022; Xie et al., 2022; Yang et al., 2022). Furthermore, genomic and proteomic profiling studies have identified significant differences between circulating EVs from BC patients and healthy controls (HC), suggesting that EVs may be a unique source of tumor biomarkers (Lee et al., 2023; Vinik et al., 2020). Plasma EVs levels were found markedly elevated in BC patients compared to HC, and that EVs levels returned to the values similar with those observed in HC following cancer surgery (Morasso et al., 2022; Vinik et al., 2020). In addition, previous proteomic analyses of EVs contents from BC patients have demonstrated good diagnostic performance, identifying potential EVs-associated proteins such as fibronectin and Del1 as possible biomarkers for BC (Lee et al., 2021; Moon et al., 2016).

Despite the encouraging results, the study of circulating EVs remains complex due to their relatively low abundance and intricate composition, which pose significant challenges for comprehensive analysis (Welsh et al., 2024). Furthermore, circulating EVs may interact with other plasma components such as circulating proteins and lipoproteins (LPs), forming what is known as the “EVs corona” (Heidarzadeh et al., 2023). The interaction between EVs and LPs, in particular, can potentially alter the biological effects of EVs on target cells (Busatto et al., 2022; Ghebosu et al., 2024).

In this context, Raman spectroscopy (RS) could be a valid approach to study the overall biochemical profile of EVs (Welsh et al., 2024). RS is a label-free technique based on the interaction between light and the molecules in a sample. It has been extensively used for the characterization of biomolecules due to its simplicity, low cost, and ability to analyze a wide range of substrates, from isolated molecules to cells and whole organisms (Butler et al., 2016; Cutshaw et al., 2024). By looking at the shift and intensity of the light scattered, RS is able to prove simultaneously multiple information on the composition and molecular structure of EVs but also of other extracellular particles (EPs), starting from an extremely limited amount of material (Bonizzi et al., 2023; Gualerzi et al., 2017; Morasso et al., 2020; O’Toole et al., 2024; Ricciardi et al., 2020).

To achieve a comprehensive overview of the biochemical profile of EVs in the plasma of BC patients, we utilized RS to first assess the differences between various classes of EPs in the blood, including different LPs (VLDL (Very Low-Density Lipoprotein), HDL (High-Density Lipoprotein) and LDL (Low-Density Lipoprotein)) and EVs of different sizes, such as large EVs (lEVs) approximately larger than 100 nm in size, and small EVs (sEVs) smaller than 100 nm. Secondly, we evaluated RS’s ability to provide reliable information on the quantities of major classes of biomolecules present comparing the results by RS with those that can be obtained using traditional biochemical assays. Lastly, we compared the compositional differences of lEVs and sEVs between BC patients and HC particularly focusing on the detection of those variation that could be due to the effect of the EVs corona.

## 2. Materials and Methods

### 2.1 Patient selection

Thirty-four early breast cancer (BC, n=34) patients and 30 healthy controls (HC, n=30), matched for sex and age were enrolled in the protocol 2490 approved by the Ethical Committee of ICS Maugeri IRCCS (Pavia, Italy) and conducted in compliance with the Declaration of Helsinki. All subjects included in this study provided written and informed consent. The main clinical characteristics of the study cohorts are summarized in **Table 1**.

**Table 1.** Demographic characteristics of the enrolled cohorts.

|  | BC | HC |
| --- | --- | --- |
| Age, years | 63,03± 12,71 | 62,43 ± 13,5 |
| BMI, Kg/m<sup>2</sup> | 24,94 ± 5,78 | 25,31 ± 5,07 |

BC: breast cancer (n=34); HC, healthy controls (n=30). Data are presented as mean ± standard deviation.

### 2.2 Blood collection

Peripheral venous blood was collected from each subject in 2 EDTA-coated vacutainer tubes. Plasma samples were obtained by blood centrifugation at 2000 ×g for 10 min at room temperature (RT). To remove cell debris and platelets, plasma samples were centrifuged at 2500 ×g for 10 min at 4°C and platelet-free plasma was stored in sterile cryovials at -80°C until use.

### 2.3 EVs isolation from human plasma samples

EVs isolation was performed through a previously reported protocol based on differential ultracentrifugation (UC) in an Optima MAX-TL ultracentrifuge (Morasso et al., 2020). Briefly, 1 ml of plasma was centrifuged at 20,000 ×g, for 1 hour, at 4°C, obtaining the lEVs in the pellet and sEVs in the supernatant. The pellet was then washed with filtered Phosphate Buffered Saline pH 7.4 (PBS 1×), centrifuged once again and the lEVs pellet was processed for downstream analysis. The supernatant from the first centrifugation was filtered through a 0.22 µm syringe filter and ultracentrifuged at 100,000 ×g, for 1 hour, at 4°C. The resulting pellet was washed with filtered PBS 1×, ultracentrifuged again in the same condition and the sEVs pellet was processed for further analysis.

### 2.4 LPs isolation from human plasma samples

The major classes of LPs (VLDL, LDL, and HDL) were isolated from human plasma using UC in a discontinuous potassium bromide (KBr) density gradient, as described by Ricciardi et al., 2020 . After the initial ultracentrifugation at 100,000 ×g for 1 hour at 4°C, VLDL and LDL were collected, while the isolated HDL fraction underwent a second UC step in the same condition to remove albumin as previously described (Ricciardi et al., 2020). Subsequently, all LPs classes were dialyzed overnight in 10 mM PBS 1× to remove KBr and stored at -80°C until use.

### 2.5 Nanoparticle Tracking Analysis (NTA)

Nanoparticle tracking analysis (NTA) was performed to detect the size and concentration of plasma-derived EVs using the NS300 instrument (NanoSight, Amesbury, UK). Samples were diluted in 1:50 ratio in filtered PBS 1× to obtain an optimal concentration of 10^8^-10^9^ particles/ml and 1 ml of the diluted sample was loaded into the instrument at a rate of 30 frames/s. Videos of particle movement were recorded three times for each sample, and the concentration and size of EVs were analysed using the NTA software (version 2.2, NanoSight).

### 2.6 Transmission Electron Microscopy (TEM) analysis

Transmission Electron Microscopy (TEM) was utilized to investigate the morphology of EPs. A suspension of 20 µl was deposited onto a carbon-coated EM grid and fixed overnight at 4°C with 25% buffered glutaraldehyde. Following fixation, the sample underwent staining with 2% uranyl acetate for 30 seconds. The samples were observed on the Tecnai Spirit 10 TEM (FEI).

### 2.7 Lipid extraction and quantification

The lipids were extracted from biological samples following the Folch procedure. Briefly, a biological sample was diluted with chloroform/methanol (2:1; v/v) and 1 ml of physiological solution (0.9% NaCl) was added. The solution was then centrifuged at 4000 rpm for 10 minutes at RT to separate the upper water/methanol phase from the lower chloroform phase. The upper phase was discarded, and the lower lipid phase was collected and stored at - 20°C until further use.

The levels of cholesterol (CL), and triacylglycerols (TG) in lipid extracts were determined using commercially available kits (Scalvo Diagnostic, #cat B74182716, #cat #B74182719) following the manufacturers’ protocols as previously described (Cazzola et al., 2013). Additionally, the phospholipid (PL) content was determined using the Bartlett assay (Cazzola et al., 2013). All measurements were conducted at least in duplicate.

### 2.8 Western blotting (WB) analysis

Protein quantification of each EPs was conducted using the Bradford assay and the absorbance was measured at 595 nm (Mechanism of dye response and interference in the Bradford protein assay). Purified EPs were resuspended in not reducing conditions Laemmli buffer for the detection of tetraspanins CD9, CD63, CD81 and apolipoproteins (Apo-A1 and Apo-B100) and boiled for 5 min at 95°C.

The lysis of EPs was performed in RIPA buffer for 20 min in ice. Following centrifugation at 18,000 × g for 20 min at 4 °C, the protein content in the supernatant was measured using the Bradford assay. EP were resuspended in reducing buffer and boiled for 5 min at 95°C. Proteins were separated on 12% SDS-PAGE polyacrylamide gels and transferred to Polyvinylidene fluoride (PVDF) membranes (Bio-Rad Laboratories). The membranes were blocked using a blocking buffer Tris-buffered saline containing 0.1% Tween-20 (TBS-T), supplemented with 5% bovine serum albumin (BSA) for1 hour at RT and then incubated with primary antibodies, anti-CD9 (1:1000, BD Pharmingen), anti-CD63 (1:1000, HansaBiomed), anti-CD81 (1:1000, Ancell), anti-Alix (1:1000, Santa Cruz Biotechnology), anti-ApoA1 (1:1000, Genetex), and anti-ApoB100 (1:1000, Abcam) at 4 °C overnight. All primary antibodies were diluted in Blocking buffer. The membranes were washed three times with TBS-T, repeated twice, and then incubated with the secondary anti-rabbit antibody conjugated with horseradish peroxidase (1:5000 cat# ab97200 Abcam) or the secondary anti-mouse antibody conjugated with horseradish peroxidase(1:5000 cat# ab97265)Abcam) for 2 hours at RT. The secondary antibodies were diluted in blocking buffer. The membranes were washed three times with TBS-T and analyzed with a ChemiDoc System (Bio-Rad Laboratories).

### 2.9 Raman spectra acquisition and preprocessing

Raman spectra were acquired using an InVia Reflex confocal Raman microscope [Renishaw, Wootton-under-Edge, UK] equipped with a solid-state laser light source operating at 785 nm.

In order to perform Raman spectral acquisition, an aliquot of 4 μl was drawn from the sample and deposited onto the surface of Raman grade CaF2 discs (Crystran, UK), followed by drying for approximately 20 minutes at RT. The Raman analysis utilized a 785-nm excitation laser with 100% power, a 1200 l/mm grating, and a 100× objective [Leica]. Spectra were acquired in the region between 670-1760 cm^-1.^ For each sample, five spectra were acquired from different positions on the drop. Cosmic rays were removed using Wire 5.5 software (Renishaw, UK). For spectra pre-processing, a Savitzky–Golay filter (parameters: Window = 7; Polynomial Order = 2) and Asymmetryc Least Square Smoothing as Asymmetrycally Reweighted (parameters: Smoothing constant = 1000 000; Max Iterations = 10; Weighting Deviations = 0.5) were used for spectral de-noising and smoothing using Orange software [PMID: 34571947]. Spectra were normalized using an Area Normalization (lower limit 1436, upper limit 1438) and the average of five spectra was considered as the representative spectrum for each subject.

### 2.10 Statistical analysis

For the statistical analysis, Kolmogorov-Smirnov and Shapiro-Wilk tests were applied to verify the normal distribution of data. Then, parametric [t test] or non-parametric [Mann-Whitney test] tests were applied as appropriate to compare mean values between the different groups using Origin software. Statistical significance was set at p-value < 0.05.

Multivariate data analysis for the automatic classification of the spectra was based on a principal component analysis [PCA] and performed using Orange software. A Support Vector Machine (SVM) algorithm was used for the analysis of RS data to discriminate between the two classes of subjects included.

## 3. Results

### 3.1 Characterization of EPs isolated from human plasma

The five main classes of EPs present in blood were isolated from human plasma of 5 BC and 5 HC using a density-gradient UC step for LPs such as VLDL, LDL and HDL, or a differential UC step for EVs, including both lEVs and sEVs. EPs were characterized according to the guidelines of the International Society for Extracellular Vesicles (*ISEV)* published in 2018 (Théry et al., 2018) and recently reaffirmed in the new MISEV2023 position paper (Welsh et al., 2024). Various techniques including NTA, TEM, and WB were employed to validate the obtained sizes, morphology, and biochemical characteristics (**Figure 1**).

**Figure 1.**
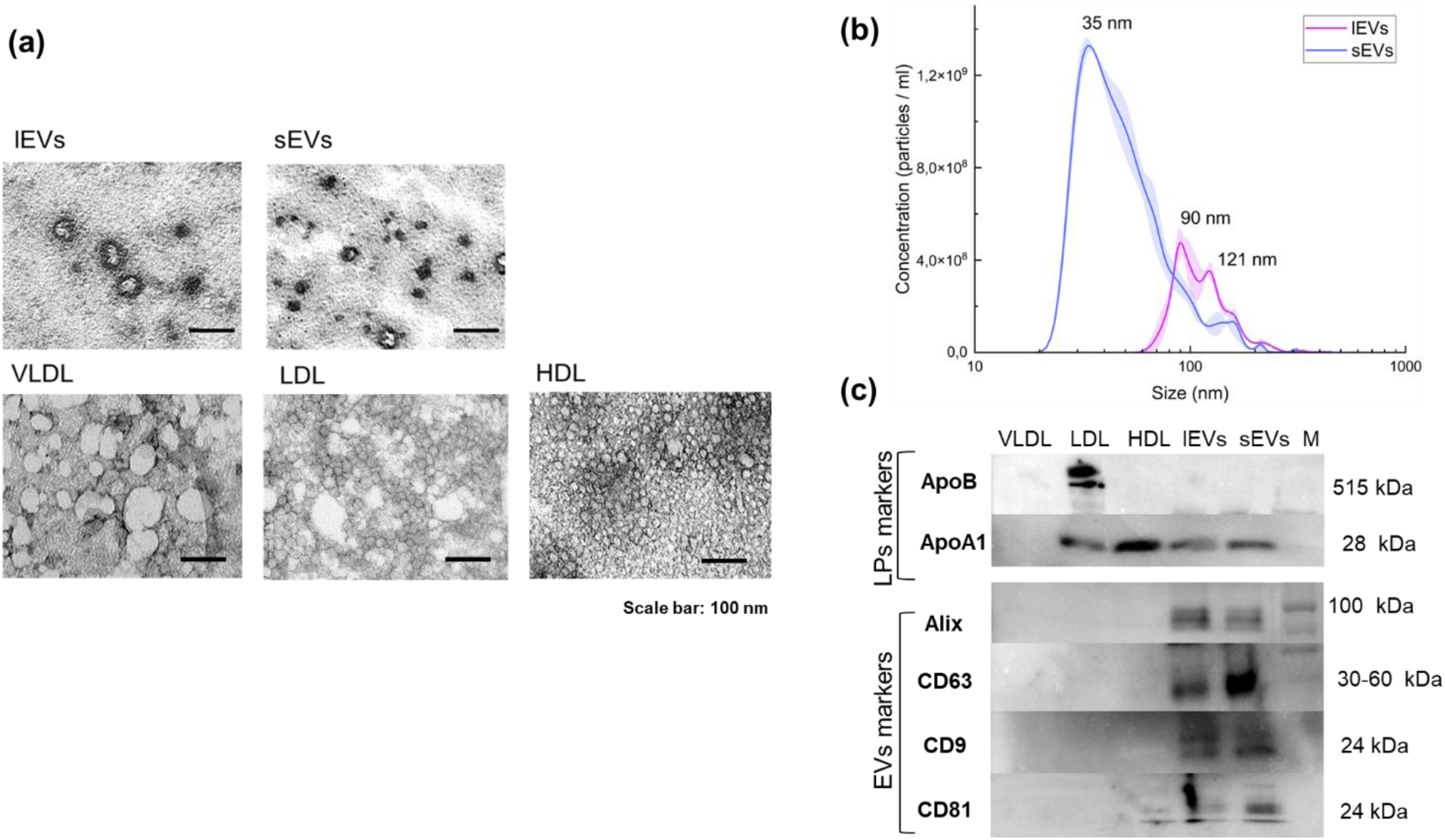
(a) TEM images of the EPs isolated from human plasma; black scale bar=100 nm **(a)**; **(b)** Size distribution and concentration of EVs quantified by NTA; **(c)** Western blot analysis of EVs markers (Alix, CD63, CD9, and CD81) and LPs markers (ApoA1 and ApoB); Molecular size (M).

TEM images revealed distinctive morphologies of the five classes of EPs obtained. EVs presented a lipid bilayer surrounding a more electrodense core. lEVs exhibited an average size of approximately 70 nm while sEVs appeared slightly smaller with a dimension of about 50 nm. LPs displayed a different dimension according to their subtype with a clear lipid core attributed to the lipids they carry **(Figure 1a)**.

The number and size distribution of isolated EVs were also quantified through NTA. sEVs presented a rather homogeneous distribution centred at 35 nm of size with a concentration of about 1.3 ± 0.02 1^0^9^ particles/ml. lEVs obtained were more polydisperse with two main populations of particles centred at 90, and 121 nm, with a concentration of 4.7 ± 0.5 10^^8^ and 3.5 ± 0.3 10^^8^ particle /ml respectively (**Figure 1b**). NTA analysis was exclusively conducted on EVs rather than LPs due to the instrument’s detection limit being insufficient to accurately capture the smaller HDL and LDL.

The presence of specific protein markers characterizing the different EPs was assessed by WB analysis, with equal amounts of sample loaded for each EPs. The obtained results confirmed the presence of the typical EVs markers CD63, CD9, CD81, and ALIX in both lEVs and sEVs samples (Nieuwland and Siljander, 2024). Small traces of CD81 were also detected in the HDL sample.

In the LPs fractions, the results confirmed the presence of the typical markers such as ApoB- 100 and ApoA-I. ApoB was mainly and distinctly expressed in the LDL fraction and was not found in any of the EVs samples. On the other hand, ApoA1 was predominantly found in the HDL fraction, with a minor band also noticeable in the EVs samples, indicating some contamination of LPs in the EVs-enriched fractions (**Figure 1c**), consistent with literature on the isolation of EVs by differential UC (Brennan et al., 2020). Not surprisingly, the ApoA1 band was also visible in the LDL samples, as previously detected in proteomic studies characterizing LPs (Lepedda et al., 2013).

None of the markers were detected in the VLDL fraction, as these EPs are predominantly composed of lipids and the amount of protein loaded when similar amounts of material are loaded for all the EPs resulted in too low a concentration to be detected by WB. Overall, these data indicate that, while some inevitable cross-contamination is present in the different samples, the various EPs, including EVs and LPs, were successfully separated from human plasma samples.

### 3.2 Raman analysis of plasma EPs

Five Raman spectra were acquired from each sample of EPs for each of the 10 subjects involved in the study. **Figure 2a** shows the overlay of the average spectrum obtained from different classes of EPs providing a fingerprint of their biochemical composition. As illustrated in **Table S1** in the supplementary information, 38 peaks common to all examined EPs were identified. Based on the scientific literature on the Raman analysis of biomolecules these peaks were assigned to different classes of biomolecules such as lipids, proteins, and nucleic acids (De Gelder et al., 2007; Movasaghi et al., 2007).

**Figure 2.**
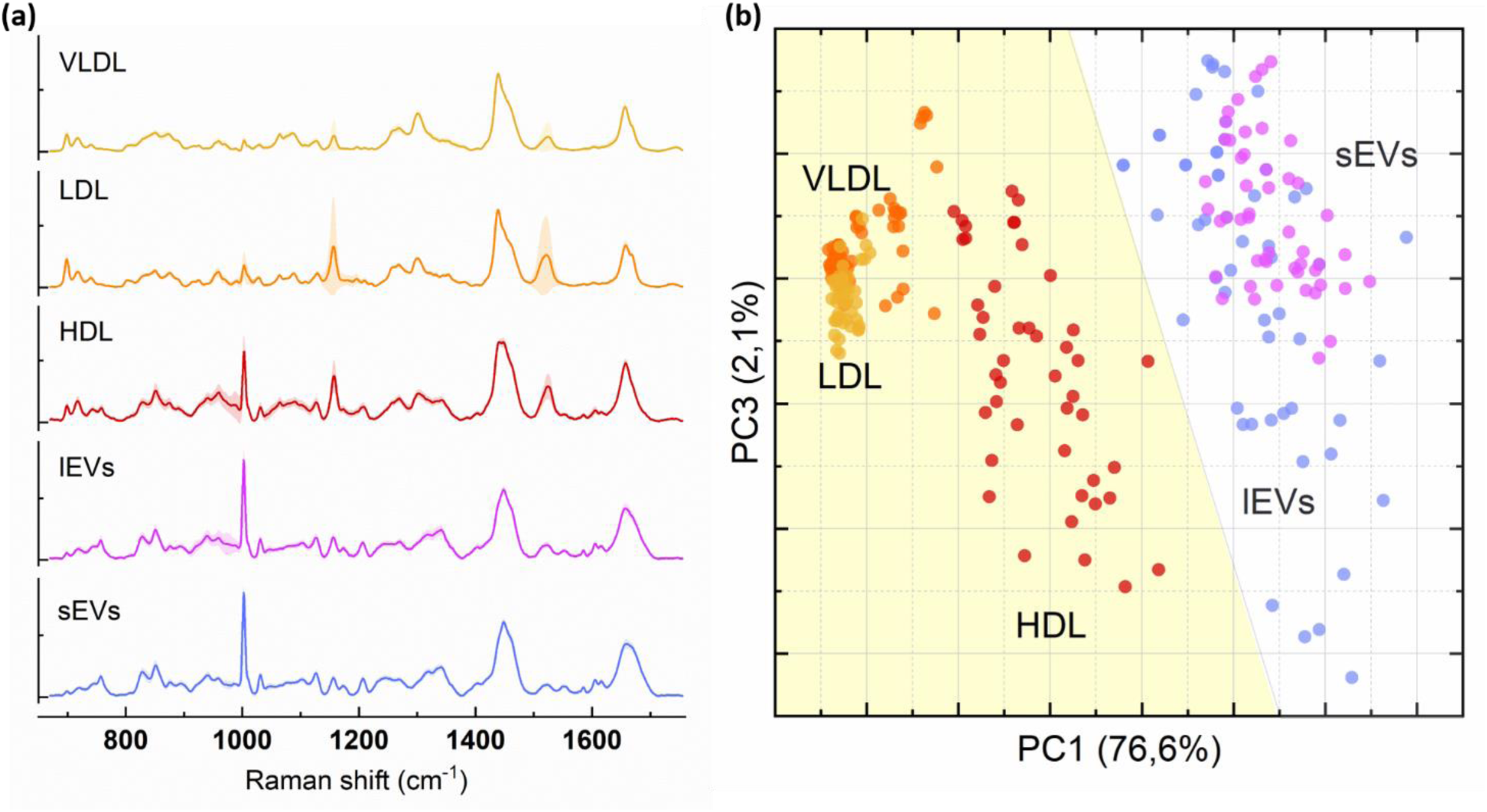
(a) Average normalised Raman spectra of the five classes of EPs (VLDL, LDL, HDL, lEVs and sEvs) from 10 subjects; **(b)** A multivariate analysis of Raman spectra based on PCA of the 38 selected features.

While the peaks appear to be consistently present in all the samples analysed, their intensity varied across the five classes of EPs, reflecting differences in their overall biochemical composition.

LPs were characterized by peaks typical of lipids, such as cholesterol (700 cm-1), phosphatidylcholine (717 cm^-1^), and carotenoids (1155 - 1520 cm^-1^), which were less intense in EVs samples. Conversely, EVs exhibited more intense peaks related to nucleic acids (746 cm^-1^ and 828 cm^-1^) and amino acids, including tryptophan (757 cm^-1^, 1339 cm^-1^ and 1552 cm^-1^) and phenylalanine (1002 cm^-1^ and 1030 cm^-1^).

A Principal Component Analysis (PCA), based on the 38 selected peaks, highlighted a clear differentiation of Raman spectra of LPs (such as HDL, LDL, and VLDL) from those of EVs (including lEVs and sEVs) (**Figure 2b**), proving the ability of RS to distinguish between the different EPs present in plasma.

However, PCA exhibited lower discriminatory power when differentiating among the subclasses of EPs. This is evidenced by the partial overlap observed between VLDL and LDL within the LP classes and between lEVs and sEVs within the EV classes, which have similar biochemical compositions.

In our dataset, the first (PC1) and third (PC3) principal components collectively described 76,6% and 2,1% of the variability respectively. The PC1 loading plot (**Figure S1a**) revealed that this differentiation was predominantly related to the lipid/protein ratio of the different EPs. Notably, for PC1, significant contributions to discrimination were observed at 700 cm^-1^ (cholesterol), 760 cm^-1^ (tryptophan), 1002 cm^-1^ (phenylalanine), 1300 cm^-1^ (lipids), 1450 cm^-1^ and 1675 cm^-1^ (proteins), and 1740 cm^-1^ (ester). On the other hand, PC3 (**Figure S1b**), which also contributed to differentiation, was characterized by peaks at 1155 cm^-1^ (carotenoids), 716 cm^-1^ (phosphatidylcholine), and 960 cm^-1^ (lipids), suggesting that this effect is mostly related to the variability in lipidic components of the EPs. Taken together, these findings offer valuable insights into the molecular signatures driving the differentiation of EPs and support the fact that RS is a powerful enough approach for the univocal identification of EVs extracted from blood plasma.

### 3.3 A comparative analysis of the biochemical composition of EPs

While it is generally accepted that the intensity of Raman peaks depends on the relative amount of the biomolecule present in the sample (Pelletier, 2003), the exact correlation between the variation recorded by Raman and the biochemical composition of EPs is not known.

We thus used a set of standard biochemical assays to quantify the level of CL, PL, TG and proteins present in each of the considered EPs and we compared the results with the correspondent Raman peaks: 700 cm^-1^ for CL, 718 cm^-1^ for PL, 1300 cm^-1^ for TG and 1246 cm^-1^ of Amide III for proteins (**Figure 3**). It is important to note that the two approaches are intrinsically different: biochemical assays detect the absolute amount of the molecule present in the sample, whereas RS provides information relative to the amount of the biomolecule in the volume illuminated by the objective. Despite this, the trends observed were remarkably similar. EVs exhibited a much higher amount of protein present both in the Raman data (**Figure 3h**) and when measured by the Bradford assay (**Figure 3d)** (Bradford, 1976). On the contrary, LPs exhibit a higher lipid content.

**Figure 3.**
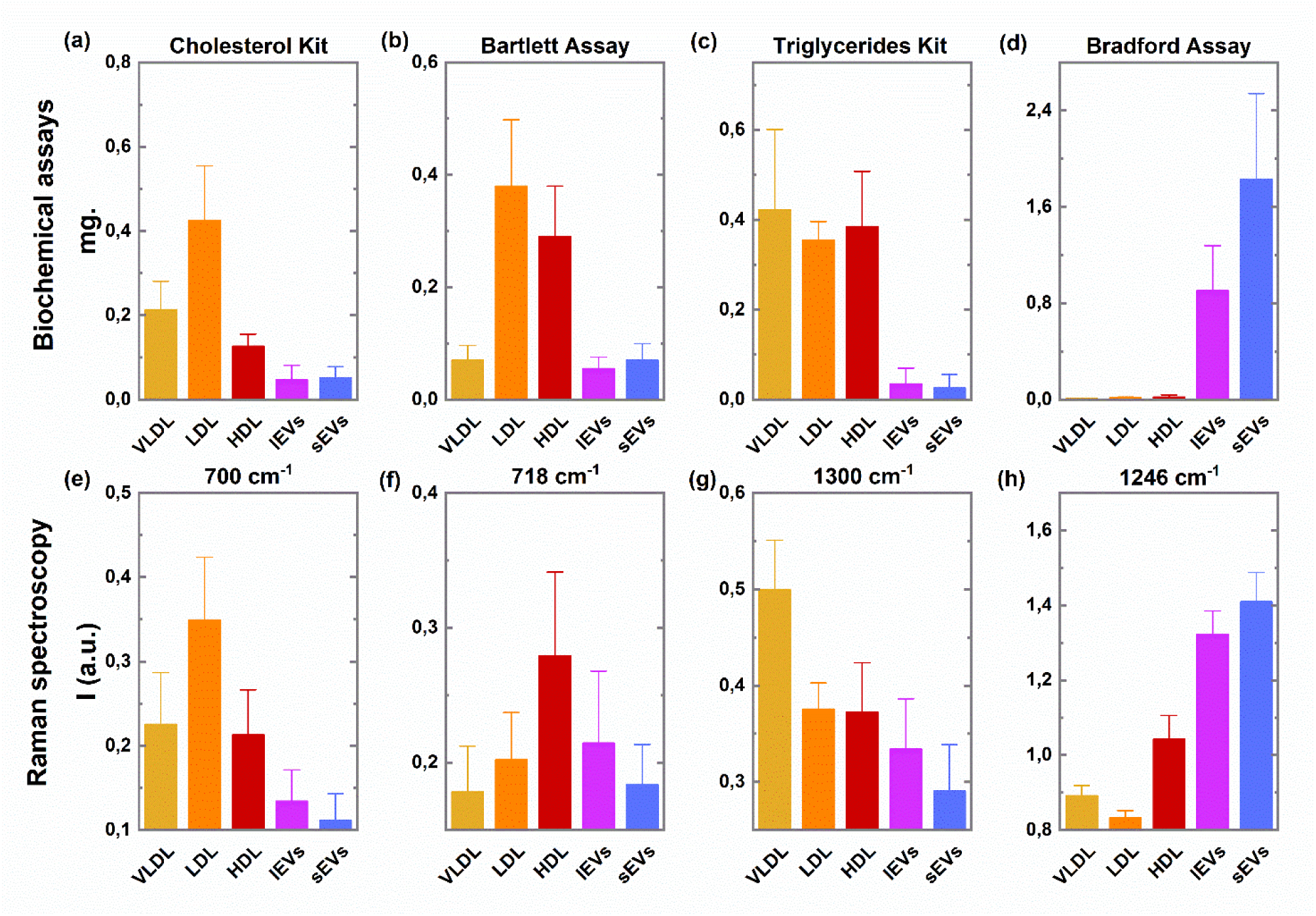
Quantification of the amount of cholesterol (CL), phospholipids (PL), triglycerides (TG) and proteins, in the different classes of EPs; Comparison between the biochemical assays **(a-d)** and the RS data **(e-h).**

Notably, CL content was enriched in the LDL fraction as evidenced by the intensity trends of the 700 cm^-1^ (**Figure 3e**) and measurements using the commercial CL dosing kit by Sclavo (**Fig. 3a**). This trend aligns well with the documented composition of LPs in the literature, where LDL is recognized as the primary transporter of CL. TG were most pronounced in the triglyceride-rich VLDL fraction reflecting its distinct role in transporting endogenous TG from the liver to peripheral tissues (Ricciardi et al., 2020). Interestingly, even though the TG dosing kit reported higher levels in VLDL (**Figure 3c**), the Raman data highlighted the difference between VLDL and other LPs classes more clearly, allowing for better discrimination of EPs (**Figure 3g**).

PL levels were significantly higher in both LDL and HDL fractions when measured using the Bartlett assay (Bartlett. 1959) (**Figure 3b**). In this case, RS data did not fully correlate (**Figure 3f**). Specifically, Raman data indicated increased PL levels primarily in the HDL fraction, known to be smaller and thus characterized by a high surface-to-volume ratio (Massey and Pownall, 1998). The differences observed between the two methods could be attributed to their targeting of different functional groups. Bartlett assay quantity phosphorous in the extracted lipids; the 718 cm^-1^ peak is instead characteristic of the C- NH3^+^ group of choline and thus refers only to the class of phosphatidylcholines. As such the two approaches do not fully align. Even in this case, the Raman data align well with the known composition of HDL that are made of 50% of proteins and 25% of PL (Gordon et al., 2011).

Overall, these results confirm the ability of RS to provide structural information on the composition of EPs in a reliable and much faster manner than traditional biochemical assays, offering multiple information in a single measurement without requiring any further extraction or processing steps.

### 3.4 Raman analysis of plasma EVs in BC cancer patients

After the assessment of the ability of RS to discriminate EPs and its ability to effectively provide information on their biochemical composition, we investigated by RS the biochemical differences characteristics of EVs extracted from the plasma of BC patients. For this study, lEVs and sEVs were isolated from the plasma of 34 BC patients and 30 HC subjects. Statistical analysis of 38 peaks previously identified, is visually represented through volcano plots (**Figure 4**) and detailed in **Supplementary Tables 2 and 3**. The peaks selected as significantly diverging between the studied population had a t-test value below 0.05 and a difference between the means of the two populations of at least 5% over the overall mean of all subjects included.

**Figure 4:**
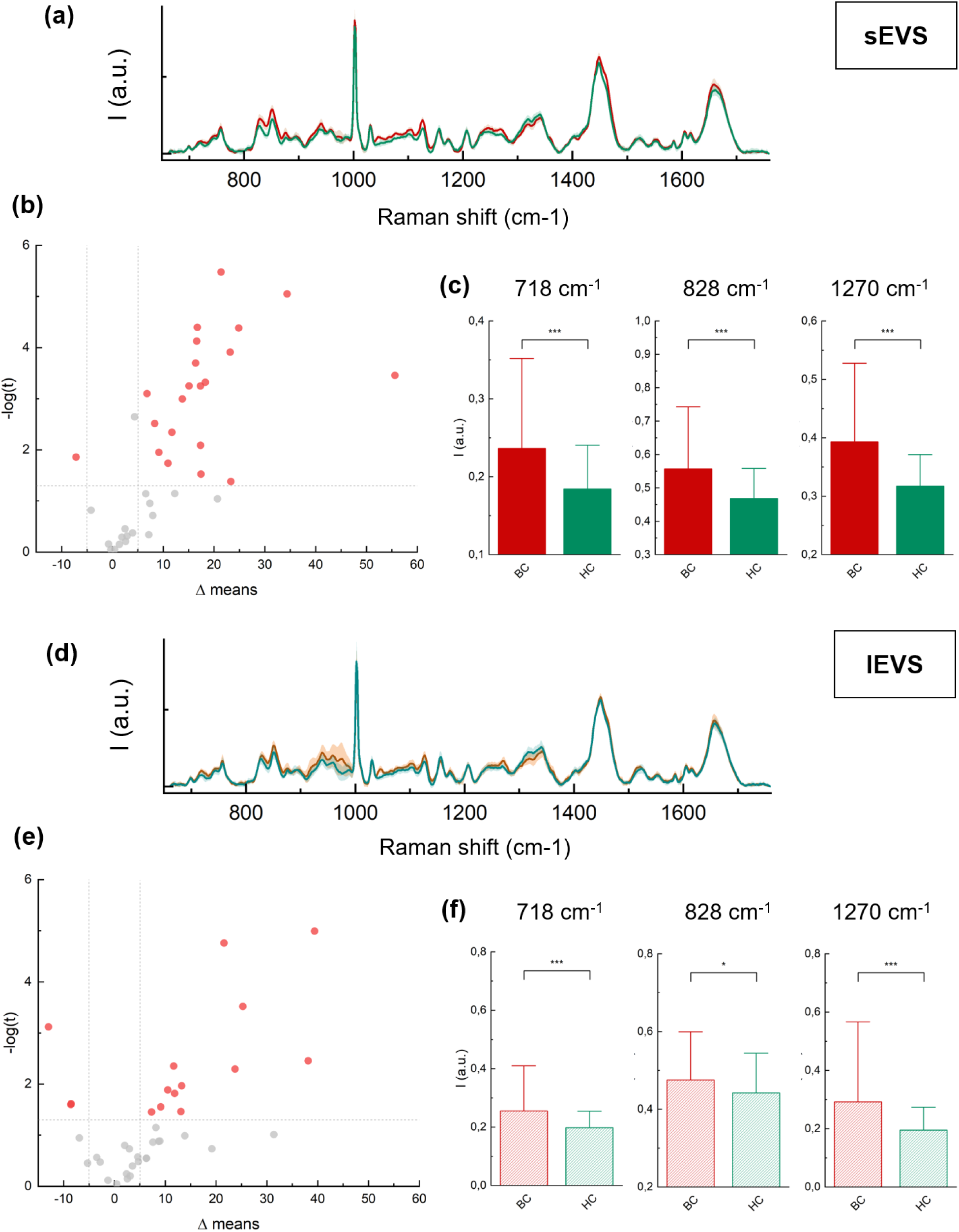
Overlay of the normalized average Raman spectra for BC and HC groups in sEVs **(a)** and lEVs **(d)**; Volcano plots illustrating statistical analysis results: 22 significant peaks in sEVs **(b)** and 15 in lEVs **(e)** from a total of 38 peaks identified in both EV populations; Box plots showing the different Raman intensity of selected peaks in BC patients vs HC subjects (c-f).

The results of the statistical analysis highlighted that the composition of sEVs was mostly affected by the presence of BC, with 22 peaks showing statistically significant differences.

In contrast, only 15 peaks were identified as significantly different between BC patients and HC in lEVs **(Figure 4b and 4e).**

The qualitative analysis of the spectral differences between the BC and HC groups is illustrated in **Figure 4a and 4d**, which display the overlay of the normalized average Raman spectra for each group across the two classes. The peaks identified as significantly different between BC and HC were primarily associated with components related to nucleic acids, and lipids, which were higher in BC samples for both sEVs and lEVs.

In particular, the most notable differences were found on both classes on peaks related to phosphatidylcholine (718 cm⁻¹), phosphodiester bonds (828 cm⁻¹), and lipids backbone (1270 cm⁻¹) as reported in the box-plot **Figure 4c and 4f**. Among the differences specifically identified on sEVs it is notable to observe that many occurs on lipidic peaks such as the peak at 1448 cm^-1^ and the one at 1656 cm^-1^. **Figure S2** reports a superposition of the mean spectra of sEVs of BC and HC with the mean spectra of LDL. Common features emerged between LDL and sEVs from BC patients, confirming an alteration in the lipid components of blood-derived sEVs in the presence of the disease. This suggests that these changes might be due to an increased interaction between LDL and sEVs in BC patients.

Overall, these data support the existence of a distinct biomolecular profile in EVs from BC patients, with particularly pronounced differences observed in sEVs. These results underscore the potential of RS in identifying a BC-specific spectral signature, highlighting its value for both diagnosis and the characterization of EPs.

### 3.5 Classification of BC and HC based on plasmatic levels of sEVs

The diagnostic potential of RS analysis of EVs was tested using a machine learning approach (Mahmoud et al., 2023) on data obtained from sEVs, as this class exhibited the most pronounced differences among groups.

A SVM algorithm with a linear kernel was developed using the 22 selected features to discriminate between the two classes of subjects included. SVM was chosen as represent a powerful supervised learning method capable of transforming high-dimensional data for binary classification. SVM has been extensively used for the analysis of RS data (Houhou and Bocklitz, 2021) as it is considered particularly appropriate as does not require that all classes analyse presented similar covariance (Guo et al., 2021). As the numerosity of the patients included in this study was too low for fully validating the results on an independent dataset, results were cross-validated by a Leave-one-patient-out (LOOCV) method. The discrimination ability of the approach was evaluated using a ROC curve analysis (**Figure 5**). The results obtained showed an AUC of O.872; Sensitivity 0.79; Specificity 0.90; PPV 0.88 and NPV 0.81. As such the Raman assay presented a very high PPV that can be of interest for the application of a liquid biopsy approaches in those area where the mammography screening cannot be implemented.

**Figure 5:**
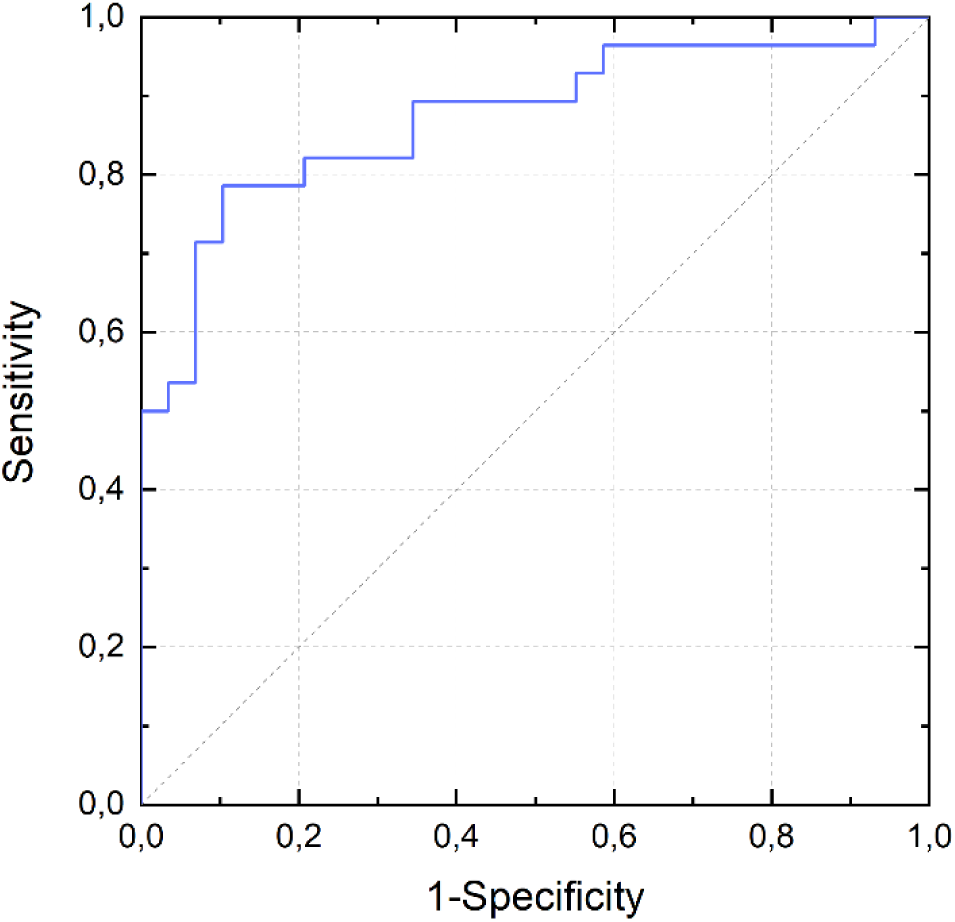
ROC curve obtained using a SVM algorithm with a linear kernel for the assignment of the Raman spectra of sEVs. Target class: BC.

## 4. Discussion

The study of EVs as potential circulating biomarkers for BC is a new potential liquid biopsy approach that could have several benefits over others that standard targets such as CTCs and nucleic acids finally allowing a precise monitoring of BC patients other all the decors of the disease.

The data on the overall variations that a pathologic state an in particular BC induce on circulating EVs however are still very limited because of the need to use a specific analytical method targeting each class of biomolecules contained in EVs.

The use of vibrational spectroscopies such as RS and IR spectroscopy to obtain a global overview of the biochemical composition of EVs that might be of diagnostic interest has been at the centre of multiple efforts in the EVs community exactly because of their operative advantages over other omics approaches such as the fast time of analysis, low cost and ability to target different classes of biomolecules in a single assay (Carlomagno et al., 2021; Gualerzi et al., 2019; Koster et al., 2022). These promises were also recently recognised in the latest version of the MISEV guidelines from 2023 where for the first time a whole paragraph was dedicated to RS reporting specific indication for the use of this approach in EVs analysis (Welsh et al., 2024). In this work, we have focused on the Raman analysis of EVs from blood as well as of other EPs such as LPs. We first studied the spectra obtained from different classes of LPs (VLDL, LDL and HDL) and from lEVs and sEVs in order to check how the spectra obtained are related with the different EPs present in blood.

EVs were obtained by differential UC, while LPs were isolated using density gradient UC in KBr. Differential ultracentrifugation was chosen for EVs because, although sometimes considered less effective than methods like size exclusion chromatography or the combination of multiple methods, it avoids the introduction of external contaminants such as PEG, glycerine, or other stabilizers, which were detected by RS when tested in our laboratory. While KBr can be effectively removed from samples by extensive dialysis, it tends to co-isolate HDL and EVs (Yuana et al., 2014), as observed in our characterization of HDL via WB. Thus, density gradient UC in KBr is an effective method for isolating HDL, which are at least three orders of magnitude more abundant than EVs, but it is not suitable for isolating EVs (Simonsen, 2017). While hints of co-isolation of different particles were present in our samples as expected, the physical and biochemical characterization of the different classes of biomolecules in each sample supports the effective obtainment of reasonably pure fractions of EPs.

Our results showed that consisted Raman bands can be identified in all the main classes of circulating EPs but that the intensity of these bands could be used for their discrimination, a result that paves the way for the use of RS as a tool for the potential assessment of the purity of blood-derived EVs similarly to what was proposed for cell culture-derived EVs (Gualerzi et al., 2019). The results of the multivariate data analysis in particular showed how the ratio between lipids and protein present (PC1) and variations in the lipidic and generally hydrophobic components are (PC3) the driving force between the ability of RS to clearly differentiate between EVs and LPs. However, due to their similar biochemical composition, RS proved less effective in distinguishing between lEVs and sEVs. lEVs obtained by differential UC, in particular, are notably heterogeneous in their spectral profiles, as also previously observed (Morasso et al., 2020), and do not form a defined cluster in the PCA. A similar situation was found for HDL. On the contrary, sEVs and LDL were characterized by very homogeneous spectra.

RS profiles were thus validated by comparing the behaviour of some characteristics peaks with the results that can be obtained using standard biochemical assays for the quantification of CL, TG, PL, and proteins. The results obtained from the biochemical assays showed good agreement with the RS data, confirming its ability to provide structural information on the composition of EPs more quickly and at a lower cost than traditional assays for all biomolecules studied, except for PL. Differences in the behaviour of PL are due to the fact that RS targets a specific subtype of these lipids. Specifically, it detects only phosphatidylcholines, which contain a quaternary amine group with a distinct Raman shift at 718 cm^-1^ (Bik et al., 2022). On the contrary Bartlett assays quantify the phosphate groups present in the hydrophobic fractions extracted from EPs. It should be noted that RS provides clearer discrimination of subpopulations of EVs and LPs compared to biochemical assays, even though a complete separation of the classes was not achieved in the PCA. For example, RS better highlighted that VLDL are much richer in TG compared to LDL and HDL. Overall our data provide a strong rationale at the use of RS to study the biochemical composition of circulating EPs in a fast and reliable way.

Starting from these premises, we investigated RS’s ability to distinguish between EVs isolated from the plasma of HC and BC patients by studying their spectral characteristics in lEVs and sEVs. The 38 previously identified spectral regions were analyzed to determine which of these regions showed significant spectral differences between BC and HC. Differently from what observed in a previous study on Amyotrophic Lateral Sclerosis (ALS) patients where variations among classes were observed on lEVs on the peaks relative to aromatic amino acids (Morasso et al., 2020) , in BC patients the most significant differences were found on sEVs (22 out of 38 peaks). Analysing the data, the differences primarily relate to BC refer to nucleic acids (828 cm^-1^), lipids (718 cm^-1^ and 1270 cm^-1^) that were more abundant in cancer patients.

The high expression levels of nucleic acids in BC samples correlate with the uncontrolled proliferation of tumor cells, leading to increased levels of tumor-derived EVs with an altered nucleic acid cargo ((Loric et al., 2023; Rahbarghazi et al., 2019; Tai et al., 2019). Variations in the level of lipids in EVs are open to multiple possible interpretations. Firstly, this result can be attributed to variations in the double-layer structure surrounding EVs. The notable increase in lipids in BC patients can also be explained by an augmented interaction between sEVs and LDL in these patients. This hypothesis is directly supported by the comparison of the mean spectra of sEVs from HC and BC patients with the average spectra of LDL, as shown in **Figure S2**. In this figure, BC patients clearly exhibit LDL features superimposed on the sEVs spectrum. Busatto and co-workers (Busatto et al., 2020) previously demonstrated that EVs released by a BC cell line prone to metastasize to the brain specifically interact with LDL. This interaction impacts the ability of EVs to be taken up by monocytes and can occur both under mixing and in physiological conditions (Busatto et al., 2022; Iannotta et al., 2024; Sódar et al., 2016). Our data thus might represent a first hint of the relevance of the interaction between EVs and LDL in cancer, impacting lipid metabolism. Lipid metabolism is know as a critical pathway in tumor progression, which depend on substantial lipid reserves for membrane synthesis, energy production, and signalling (Llaverias et al., 2011). The interplay between EVs and LDL may enhance the lipid supply to tumor cells, thereby promoting aggressive behaviors such as proliferation, invasion, and metastasis (Li et al., 2023; Liu et al., 2022; Vogel et al., 2024).

Although the present study is limited by the number of subjects included in the analysis and all results will require further validation on a larger cohort of BC patients and HC, we performed a preliminary assessment of the diagnostic potential of RS analysis of EVs in identifying BC patients. Machine learning algorithms are particularly suitable for the analysis of numeric data such as the one provided by RS (Mahmoud et al., 2023). Here, we tested a SVM algorithm for its potential to correctly classify patients. The positive results obtained with a classification accuracy of about 85% and an AUC of 0.87 support the further exploration of the Raman analysis of EVs as a diagnostic tool for BC. This result aligns with findings from other studies on the diagnostic potential of EVs in BC (Lan et al., 2024; Shi et al., 2024; Xu et al., 2024). However, RS has the notable advantage of being reagent less and requires no further procedures beyond the extraction of EVs from plasma. This enables RS to maintain very low analytical costs (Hanna et al., 2022).

## ACKNOWLEDGEMENTS

The work was funded by the Ministry of Foreign Affairs and International Cooperation and by the Italian Ministry of Health through the project: “Identification of a specific metabolic fingerprint of breast cancer associated extracellular vesicles by Raman Spectroscopy (BrERa)” within the framework of the “project of great relevance” in the bilateral collaboration between Italy and India.

We want to thank the Institutional BioBank “Bruno Boerci” for the support in the storage and handling of the clinical samples analysed. We extend our deepest gratitude to all the patients who participated in this study.

## ETHICS APPROVAL STATEMENT

This study was approved by the Ethical Committee of ICS Maugeri IRCCS (Pavia, Italy) (protocol 2490).

